# The mechanosensory DEG/ENaC channel DEGT-1 is a proprioceptor of *C. elegans* foregut movement

**DOI:** 10.1101/2025.01.01.631014

**Authors:** Emily A. Bayer, Susan E. Mango, Oliver Hobert, Alexander F. Schier

## Abstract

The gastrointestinal tract is subjected to extensive mechanosensory stimulation during food ingestion. However, the identities of mechanosensory receptors in the enteric nervous system are largely unknown. The pharynx of *C. elegans* is a structurally and functionally homologous model of the vertebrate foregut, but is comprised of only 20 neurons that are embedded within the muscles and epithelial cells of the organ. Here we report that the DEG/ENaC family ion channel DEGT-1 is a proprioceptor of pharynx movement. DEGT-1 protein is expressed in 4 pharyngeal neurons (MI, M3, I4, and M5) and localized to their neuronal soma in direct contact with the collagenous pharyngeal basement membrane. *degt-1* mutants display abnormally rapid feeding in the presence of food, causing global changes in lipid accumulation. *degt-1* mutants also pump rapidly when pumping is induced by the presence of serotonin alone, suggesting that DEGT-1 is required for proprioception of pharyngeal pumping itself, rather than sensing ingested food. DEGT-1 is required in only two pharyngeal neurons (I4 and M5) to control pumping rate. Taken together, these results suggest that DEGT-1 modulates pharyngeal pumping rate by relaying proprioceptive feedback generated by the shear force of the pharynx against its own basement membrane. Thus, mechanosensors in enteric nervous systems modulate organ function not only by detecting forces from ingested contents, but also the movements of the organ itself.

## INTRODUCTION

One of the most fundamental processes for growth and survival of an organism is the ability to ingest and digest food. This process is facilitated by the gastrointestinal tract, which is a tube formed by laminated muscle layers. These muscle layers each have specialized properties to facilitate the digestion and passage of ingested contents. Neurons and neural processes are embedded within and between the laminar muscle layers, which contribute to motor control of digestion, but also enable sensory feedback from the digestive system. In particular, nearly a quarter of neurons along the entire length of the mammalian gastrointestinal tract encode mechanosensory information^1^. These neurons do not only include canonical sensory neurons, but also apparent inter- and motorneurons that respond to mechanical distention. It has been observed that some myenteric neurons respond to distention specifically of their neurites, while some respond to distention of the somata itself^2^. These observations suggest that in addition to considering gene expression, the differential protein localization of mechanosensory receptors within neurons may be highly relevant for the physiological function of the cell. Gene expression studies alone have been insufficient to definitively link mechanoreceptors to the functionality of enteric neurons. For instance, Piezo1 expression is abundant in enteric neurons, but it is still unclear whether it is a primary mechanoreceptor^3^, and detailed functional studies of TRP family channels has drawn into question whether they all function as mechanoreceptors^4^. While particular processes like peristalsis are known to be coordinated by mechanoreceptive neurons, the responsible mechanosensors are molecularly unknown^5^.

One family of channels that has been definitively linked to mechanosensation is the DEG/ENaC superfamily (including the proton-gated ASIC subfamily most common in mammals). DEG/ENaC channels can respond to a variety of stimuli, but in the case of mechanosensation can be tethered between a neuronal process and surrounding basement membrane, thus allowing for channel opening upon mechanical distension of the neuron^6^. DEG/ENaC/ASIC channels have been implicated specifically in enteric mechanosensation in *C. elegans, Drosophila*, and in mammals. In *C. elegans*, the DEL-3 and DEL-7 ASIC channels respond directly to bacteria in the pharyngeal lumen to regulate exploration (a satiety-related behavior)^7^. In *Drosophila*, the *PPK1* DEG/ENaC channel regulates food consumption by sensing physical fullness in posterior enteric neurons^8^. In mice, loss of individual ASIC channels can decrease, but surprisingly also increase, mechanosensitivity in the digestive system^9^. Thus, DEG/ENaC channels are promising candidates for enteric mechanosensors governing a broad variety of internal sensation and digestive regulation, for which the molecular identity of receptors is yet unknown.

The *C. elegans* digestive system is a powerful model to study the mechanisms of gastrointestinal development and function. It consists of a specialized epithelial tube allowing ingestion and digestion of bacterial food, and is made up of a foregut, midgut, and hindgut. The foregut, or pharynx, is a muscular organ in the head of the animal that uses a series of highly coordinated muscle movements to move ingested food posteriorly^10^. Like the vertebrate gut, the entire pharynx is wrapped in a collagenous basement membrane^11^. Embedded within the organ is an autonomous nervous system comprised of 20 neurons belonging to 14 distinct classes (**Fig 1A**). The processes of these neurons converge into two ring-like plexuses in the anterior and posterior pharyngeal bulbs, where they form a densely connected neuronal network^10,12^(**Fig 1B**). These neurons are vital for proper coordination of the myogenic movements underlying feeding and digestion. For instance, ablation of the M4 motor neuron is lethal due to the inability to move food from the pharynx into the intestine, whereas ablation of the I2 interneurons results in reflux of ingested food (known as the “slipping” phenotype)^13,14^. Some neurons are also sensitive to particular environmental contexts; the role of the NSM neuron in modulating locomotion speed in response to bacteria becomes apparent only when animals are transitioning from non-feeding to feeding states^7,15^. However, despite decades of study, 5 of the 14 pharyngeal neuron classes still have no known function, or have been proposed to act redundantly^12^(**Fig 1B**).

**Figure 1:**
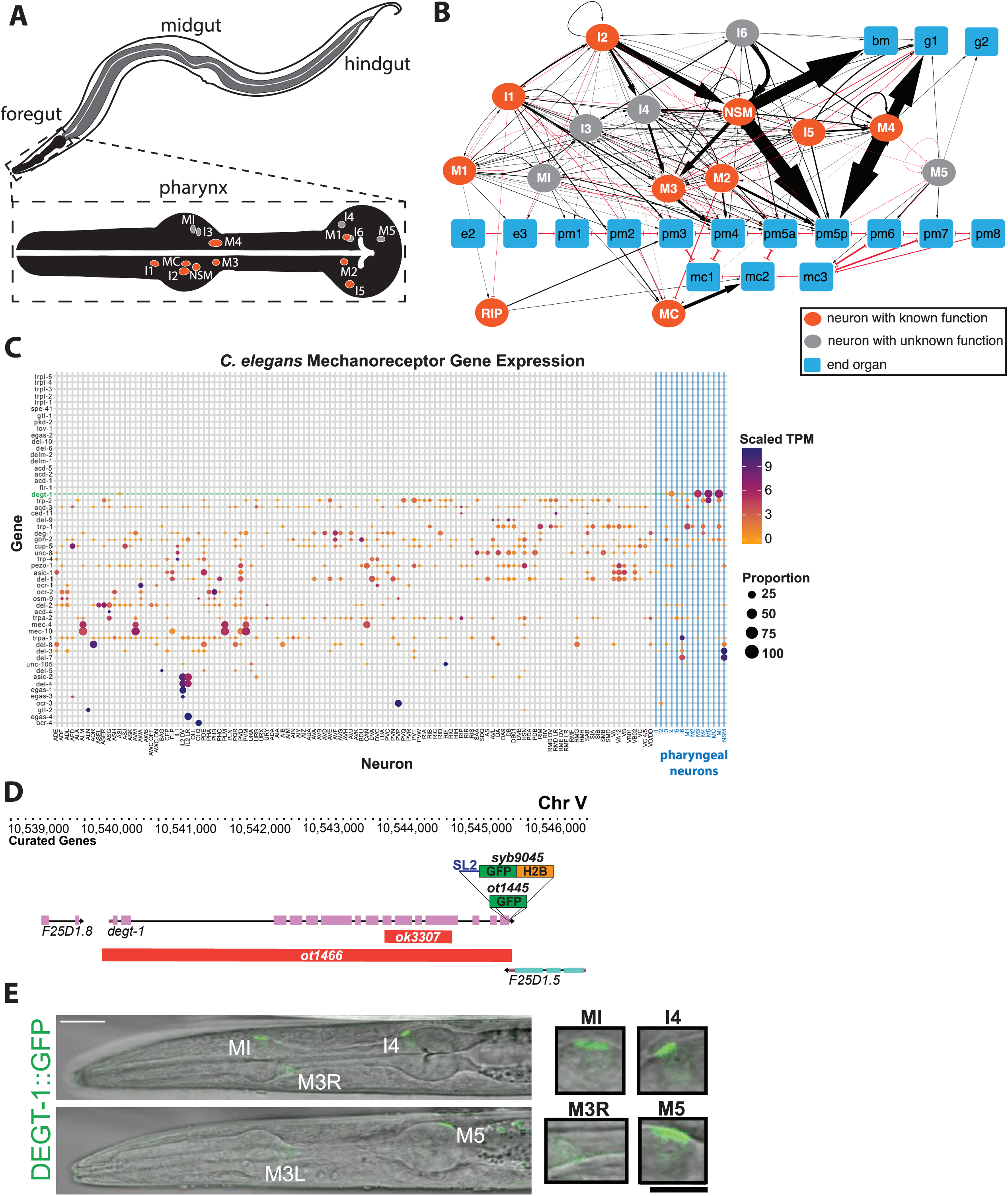
The DEG/ENaC channel DEGT-1 is expressed in four pharyngeal neuron types with previously unknown functions. **A:** Schematic of the digestive system of *C. elegans*. The magnified inset shows the position and identity of the pharyngeal neuron cell bodies within the muscle of the pharynx/foregut. Neurons with known function (summarized in ^12^) are shown in orange, neurons with unknown function are shown in grey. **B:** Connectivity of the pharyngeal nervous system and end organs (adapted from ^12^). Arrow width represents strength of synaptic connection, chemical synapses are represented by black arrowheads and electrical synapses are represented by red bracketed connectors. End organs (muscles, glands, epithelial cells, marginal cells) are depicted as blue rectangles, neurons are depicted as ovals. Neurons with known function (summarized in ^12^) are shown in orange, neurons with unknown function are shown in grey. **C:** Expression of *C. elegans* mechanoreceptors (DEG/ENaC channels, TRP channels, and Piezo, list from ^44^) across the entire nervous system based on single cell RNA sequencing dataset^16^. Each row is a gene, and each column is a neuron class. *degt-1* expression is highlighted in green, the pharyngeal nervous system is highlighted in blue. **D:** Schematic of the *degt-1* locus. Two deletion alleles of *degt-1* are shown in red bars below(*ok3307*^45^ and *ot1466*, this work), and endogenous GFP tags are schematized above (*syb9405* transcriptional reporter, *ot1445* translational reporter, both this work). **E:** DEGT-1::GFP is expressed in the MI, M3, I4, and M5 neurons. Composite GFP/DIC images of two single Z-slices of an L1 animal are shown to emphasize protein localization pattern relative to the pharynx on the left, and higher magnification images of each of the four DEGT-1+ neuron classes. White scale bar is 10uM on the left, and black scale bar for the inset images on the right is 5uM.

Here we leverage the high-resolution molecular data available for the *C. elegans* nervous system to identify a mechanosensory DEG/ENaC channel, *degt-1*, that is exclusive to the pharyngeal nervous system. We find that the pharynx senses its own muscle movements via *degt-1* to regulate feeding rate. This function is performed by two neurons of the pharyngeal nervous system, I4 and M5, that do not have previously known functions, revealing a new proprioceptive feedback module within the *C. elegans* pharynx. We demonstrate that in addition to described roles for DEG/ENaC channels in detecting the quantity and quality of ingested contents, proprioceptive sensation from the feeding organ itself is sensed to properly regulate ingestion and body size.

## RESULTS

We sought to identify mechanosensory channels that may mediate sensation of digestive muscle movements. Recent reexamination of the *C. elegans* pharyngeal nervous system connectome suggested that most pharyngeal neurons are likely sensory-motor, including many with putative sensory endings embedded in the muscle itself^12^. While sensory endings exposed to the pharyngeal lumen are capable of directly sensing ingested food, sensory endings within the muscle may rather respond to mechanical forces within the organ itself. By examining the expression patterns of putative mechanoreceptors (DEG/ENaC, TRP, Piezo) in an existing single cell RNA sequencing dataset of the *C. elegans* nervous system (CeNGEN)^16^, we noted that transcripts of the DEG/ENaC channel *degt-1* are found exclusively in the pharyngeal nervous system (**Fig 1C**). Through CRISPR/Cas9-mediated insertion of a GFP protein in the endogenous locus, we found that the DEGT-1 fusion protein is exclusively expressed in the four pharyngeal neurons MI, M3, I4, and M5 (**Figs 1C-E**). We also generated an endogenous transcriptional reporter for *degt-1* (*degt-1::SL2::gfp::H2B;* **Fig 1D**). Once again, we observed expression in MI, M3, I4, and M5, consistent with the CeNGEN data (**Supplemental Figure 1A,B**).

Several DEG/ENaC channels are known to be expressed in the *C. elegans* nervous system, such as the complex formed by the MEC-4/10 proteins and required for touch sensation^17-19^. These proteins localize to puncta along the axons of touch receptor neurons^20^. Surprisingly, using our translational reporter, we found that DEGT-1 protein does not localize to the neurites of pharyngeal neurons, but rather to the soma of the four neurons (**Fig. 1E**). In all four of these cases, the surface localizing DEGT-1 protein was directly at the edge of the pharynx. The *C. elegans* pharynx is wrapped in a collagen-rich extracellular basement membrane^11^, and DEG/ENaC channels have been previously described to associate with ECM collagens^21-23^. At the ultrastructural level, pharyngeal neurons including M5 have also been observed to directly contact the pharyngeal basement membrane^10^. We visualized DEGT-1::GFP protein alongside EMB-9/α1 collagen IV::mCherry, and found that DEGT-1 colocalizes with the pharyngeal basement membrane (**Fig. 2A**). This suggests that DEGT-1 is indeed interacting with the pharyngeal basement membrane.

**Figure 2:**
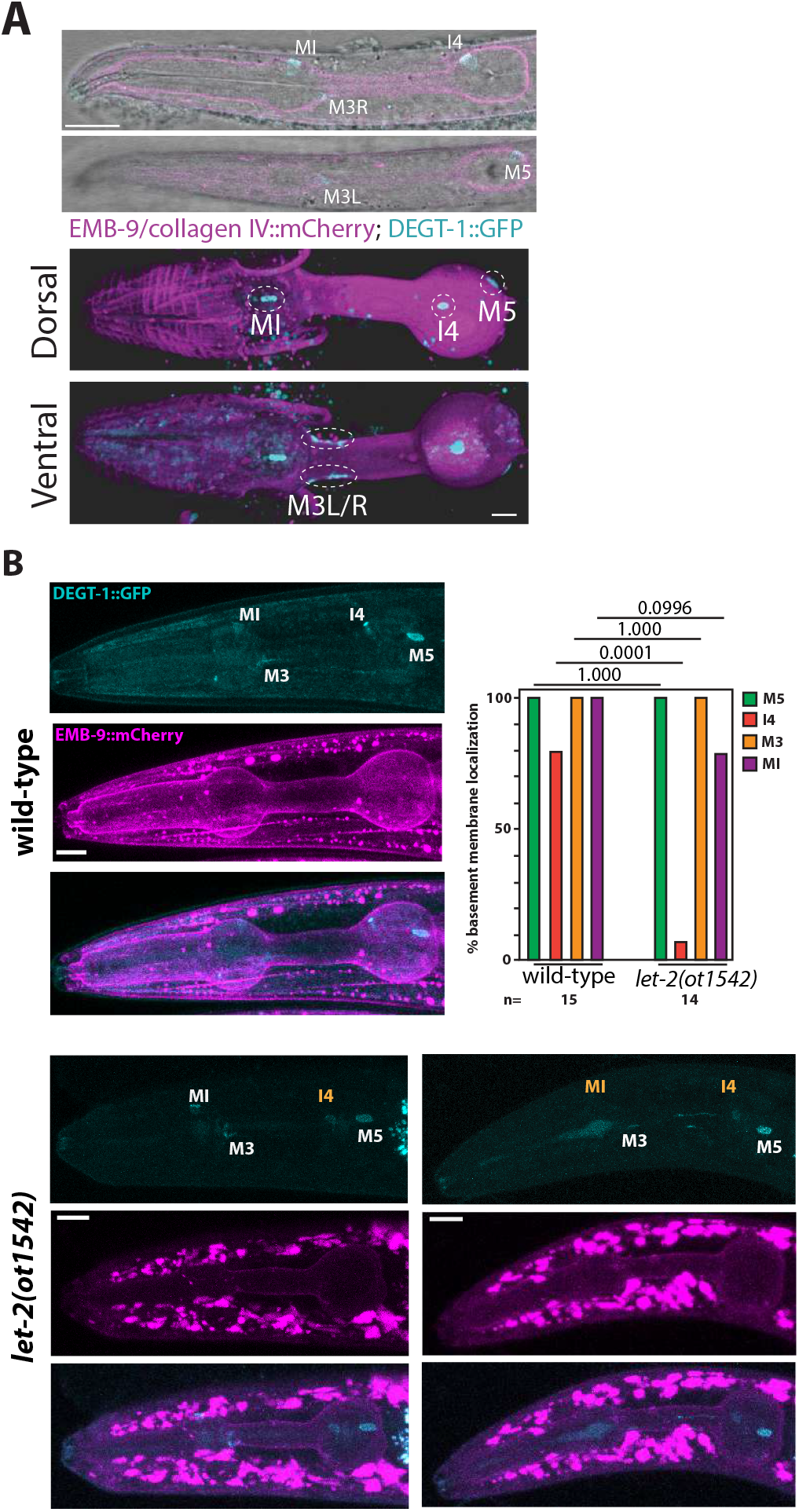
DEGT-1 localizes to neuronal cell membranes adjacent to the pharyngeal basement membrane in a collagen-dependent manner. **A:** DEGT-1::GFP colocalizes with the pharyngeal basement membrane. Above, composite DEGT-1::GFP (cyan)/EMB-9::mCherry (magenta)/DIC images in an L1 animal. Two single slices are shown to demonstrate overlap of DEGT-1 protein with collagen protein. Below, a 3D rendering of an adult animal where the pharynx was surgically exposed is shown from the dorsal and ventral sides. DEGT-1+ neurons are labeled in white on both images, scale bars are 10uM. **B:** DEGT-1 is anchored to the pharyngeal basement membrane via collagen IV. Maximum intensity projections of DEGT-1::GFP in cyan and EMB-9::mCherry in magenta, with colocalization below, of both wild-type and *let-2* mutant animals. White scale bars are 10uM. Neurons are labeled in white where they were observed to still have DEGT-1 colocalized with the basement membrane, and yellow where colocalization was lost. To display the range of phenotypes, images of two *let-2* mutant animals are shown. The plot in the upper right displays the % of animals in which each of the 4 DEGT-1+ neurons showed DEGT-1 protein colocalized with the basement membrane. p-values (above plot) calculated by Fisher’s exact test.

In the context of *C. elegans* body wall muscle, the UNC-105 DEG/ENaC channel has been found to interact both genetically and functionally with the α2 collagen IV encoded by *let*-2^22^. Both the body wall muscle and pharynx are similarly apposed by collagen-rich basement membranes^11,24^. Thus, we examined whether LET-2 is required for the localization of DEGT-1 to the pharyngeal basement membrane. The initial establishment of basement membranes is essential for completion of embryogenesis, but depleting basement membrane collagens during larval development is viable^24,25^. We utilized a temperature-sensitive mutation in *let-2* that allowed us to selectively perturb basement membrane collagen during larval development^26^, while monitoring basement membrane integrity using the EMB-9/α1 collagen IV::mCherry transgene. We observed that, similar to prior RNAi-based approaches of depleting *let-2*, there was lower fluorescence intensity of EMB-9::mCherry, ectopic accumulation of collagen in the pseudocoelomic space in the head, and frequent deformation of pharyngeal morphology, although we did not observe disruption or gaps in continuity of the basement membrane (except in rare cases where the entire pharynx tore in half; **Fig 2B**). Because the basement membrane was still continuous, we were able to test whether post-embryonic impairment of LET-2 resulted in loss of DEGT-1 tethering to the basement membrane collagen. We found differential sensitivity of DEGT-1+ neurons to loss of LET-2; DEGT-1 localization to the basement membrane was almost always lost in the I4 neuron, sometimes lost in the MI neuron, and never lost in the M3 or M5 neurons (**Fig 2B**). Under normal conditions, DEGT-1 in the I4 soma displays the smallest region of colocalization with the basement membrane (**Supplemental Fig. 2A**), suggesting that I4 may be the most susceptible to post-embryonic partial disruption of the basement membrane.

It has also been posited that DEG/ENaC channels may provide structural connection between tissues and basement membranes^22^. Thus, we tested whether DEGT-1 is required for the soma of pharyngeal neurons to colocalize with the basement membrane. We labeled the cytoplasm of all pharyngeal neurons in GFP, and quantified colocalization of DEGT-1+ neuronal soma with the basement membrane. Using a CRISPR-generated deletion of the entire *degt-1* locus (**Fig. 1D**), we found that all four normally DEGT-1+ neurons maintain their direct apposition with the pharyngeal basement membrane, even when DEGT-1 was not present (**Supplemental Fig. 2B**). Thus, loss of *degt-1* does not result in general defects in anatomy of the pharyngeal nervous system.

Because *degt-1* mutants displayed normal neuroanatomy, and DEG/ENaC channels have well-established roles in mechanosensation^18,27,28^, we hypothesized that DEGT-1 may be acting as a mechanosensory channel in pharyngeal neurons. One of the DEGT-1+ neurons, M3, acts as a proprioceptor for pharyngeal muscle contraction ^14^, and regulates the duration of pharyngeal pumping via cholinergic neurotransmission ^29^, but the mechanosensory channel responsible for primary sensation is unknown. None of the other three DEGT-1+ neurons, MI/I4/M5, have any experimentally determined function^12^. Network analysis of the connectivity of these three neuron classes has yielded predictions that each of the three neurons function in regulating different aspects of pharyngeal pumping (MI: pumping, I4: neuromodulation, M5: grinding), but these predictions have not been experimentally confirmed^12^.

We first assessed whether DEGT-1 is required to regulate pharyngeal pumping under standard laboratory conditions, with animals freely feeding on bacterial lawns. We found that in the absence of DEGT-1, the pharyngeal pumping frequency of freely feeding animals increased from ∼4.5Hz to ∼5.75Hz (**Fig. 3A**). This phenotype also allowed us to confirm that our translational DEGT-1::GFP fusion generates a functional protein, as DEGT-1::GFP animals did not display any increase in pharyngeal pumping frequency (**Supp Fig 3A**). An increase in pharyngeal pumping frequency is not only evidence of dysregulation of the function of the organ itself, but suggests that animals have an increase in food consumption and thus fat storage. To determine whether increased pumping resulted in physiological changes, we measured the length and width of *degt-1* mutants in the last larval stage (L4) and in early adulthood. We found that L4 stage animals did not differ from wild-type, but that once animals reached adulthood, they were wider, but not longer (**Supp. Fig. 3B**). By staining the fat storage compartment with the fixative-based dye Oil Red O^30^, we found that the increased width of *degt-1* mutants was accompanied by an increase in intestinal fat storage (**Fig. 3B**).

**Figure 3:**
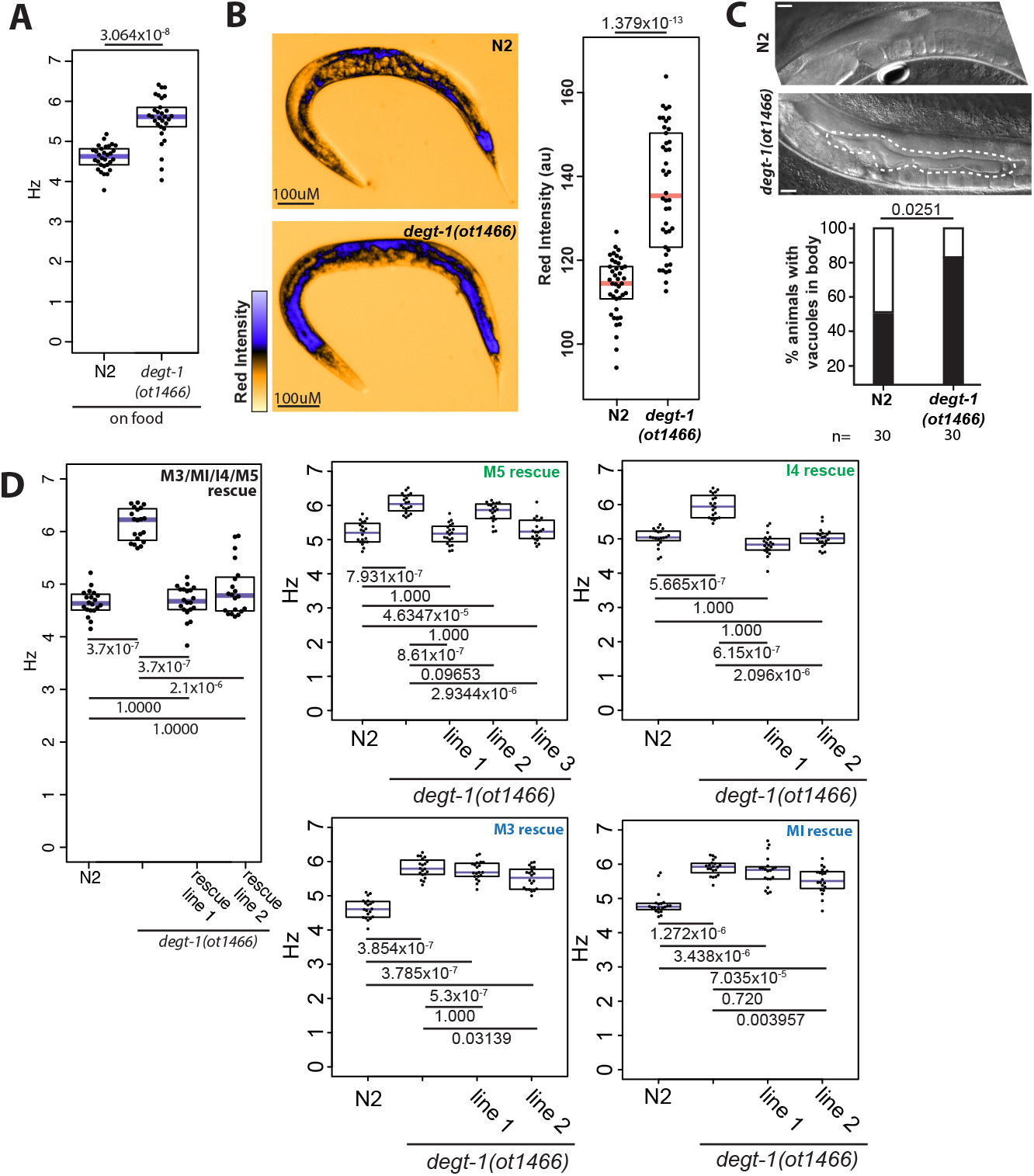
DEGT-1 controls feeding rate via the I4 and M5 neurons. **A:** *degt-1* mutants pump more rapidly than N2 (wild-type) animals when freely-feeding on bacteria. Pumping frequency is displayed in Hz, each black dot represents one animal, purple bars are medians, black boxes are quartiles. p-value (above plot) calculated by Wilcoxon rank sum test. **B:** *degt-1* mutants accumulate excess intestinal lipid. Red channel of RGB image is shown on the left (pseudocolored using the ICA LUT (Fiji)), quantification of red intensity of intestinal staining is shown on the right. Each dot represents the intestinal intensity of one animal, salmon bar is median, boxes are quartiles. p-value (shown above plot) was calculated by Wilcoxon rank-sum test. **C:** *degt-1* mutants accumulate vacuoles in the body cavity by day 5 of adulthood. While most N2 (wild-type) animals do not display vacuoles in the body cavity at day 5 (above), *degt-1* mutants show large vacuoles (encircled by dashed white line, below). Scale bars 20uM. p-value (above plot) calculated using Fisher’s exact test. **D:** Rapid pumping in *degt-1* mutants is rescued by re-expression of DEGT-1 cDNA in the I4 and M5 neurons. On the left, rescue in all four DEGT-1+ neurons is shown, and on the right, rescue in individual neurons is shown. Two independent transgenic rescuing lines are shown for each construct, with the exception of M5 rescue, where three independent lines were tested to resolve conflicting results between the first two transgenic lines. Pumping frequency is displayed in Hz, each black dot represents one animal, purple bars are medians, black boxes are quartiles. Adjusted p-values were calculated using the Wilcoxon rank-sum test with Bonferroni correction for multiple testing.

In addition to a direct link between pharyngeal pumping rate and premature death (via muscle breakdown and bacterial invasion)^31^, mutants that store excess fat have been shown to exhibit a range of accelerated aging pathologies, including reduced life span and gonad necrosis^32^. We evaluated *degt-1* mutants for both direct (pharyngeal) and indirect (fat-induced) pathologies at day 5 of adulthood, when the reproductive period has ended and senescence phenotypes begin to become apparent^33^. By feeding animals with GFP-expressing bacterial food, we were able to evaluate whether *degt-1* mutants showed increased bacterial infection of the pharyngeal muscle due to their increased feeding rate. At day 5 of adulthood (when bacterial infection of pharyngeal muscle is still rare in wild-type animals), we did not find any increase in bacterial infection of pharyngeal muscle, or other gross phenotypes of pharyngeal muscle damage (**Supp. Fig. 3C**). In contrast, we frequently observed large vacuoles in the body cavity of *degt-1* mutants by day 5 of adulthood, an aging-associated trait that was still infrequent in wild-type animals at day 5 of adulthood (**Fig. 3C**). Thus, while *degt-1* mutants do not show direct pharyngeal pathologies as a result of increased feeding rate, they do display premature senescence presumably as an indirect effect of overfeeding.

To identify the site of action for DEGT-1 function in regulating pharyngeal pumping frequency, we generated a series of heterologous constructs to express *degt-1* cDNA in the pharyngeal nervous system using highly-specific promoter fragments (**Supp. Fig. 3D**). By re-expressing *degt-1* cDNA in all four of its normally expressing neurons (M3, MI, I4, and M5) in a *degt-1* mutant, we were able to fully restore pharyngeal pumping to the wild-type rate (**Fig. 3D**), demonstrating sufficiency of DEGT-1 in M3/MI/I4/M5.

While there is precedent for single pharyngeal neurons shaping rhythmicity of pharyngeal pumping^14,34^, it has also been proposed that other groups of pharyngeal neurons serve completely redundant functions^12,35^. It was thus possible in the context of the rapid pumping phenotype of *degt-1* mutants that one, or any combination, of the four pharyngeal neurons may be causative for the phenotype. To determine the DEGT-1 site of action, we used our heterologous expression constructs to rescue DEGT-1 in individual neurons. We found that animals with DEGT-1 rescued in the M3 or MI neurons still pumped with an increased frequency of ∼5.75Hz, but that restoring DEGT-1 expression in either the I4 or M5 neurons was sufficient to completely rescue pumping frequency to the wild-type rate of ∼5 Hz (**Fig. 3D**). Interestingly, of the four neurons that express DEGT-1, two are located in the anterior pharyngeal bulb, and two are located in the posterior pharyngeal bulb (**Fig. 1E**). Our rescue results indicate that restoring DEGT-1 expression in either of the two posterior neurons is sufficient for the regulation of pumping frequency, and that the two anterior neurons seem to be dispensable for this phenotype.

Pharyngeal pumping in *C. elegans* is comprised of multiple distinct steps. To examine how loss of *degt-1* affects pharyngeal pumping at greater resolution, we used the ScreenChip microfluidic device (InVivo Biosystems), which records the electrical activity generated by pharyngeal muscle during each pump^36,37^. First, we used the ScreenChip to evaluate whether *degt-1* mutants show enhanced feeding behavior in the absence of food, as we found they did in the presence of food. We found that, like wild-type animals, *degt-1* mutants completely inhibit pumping when no feeding stimulus is present, indicating that they do not have an indiscriminate increase in pharyngeal pumping(**Fig. 4A**).

**Figure 4:**
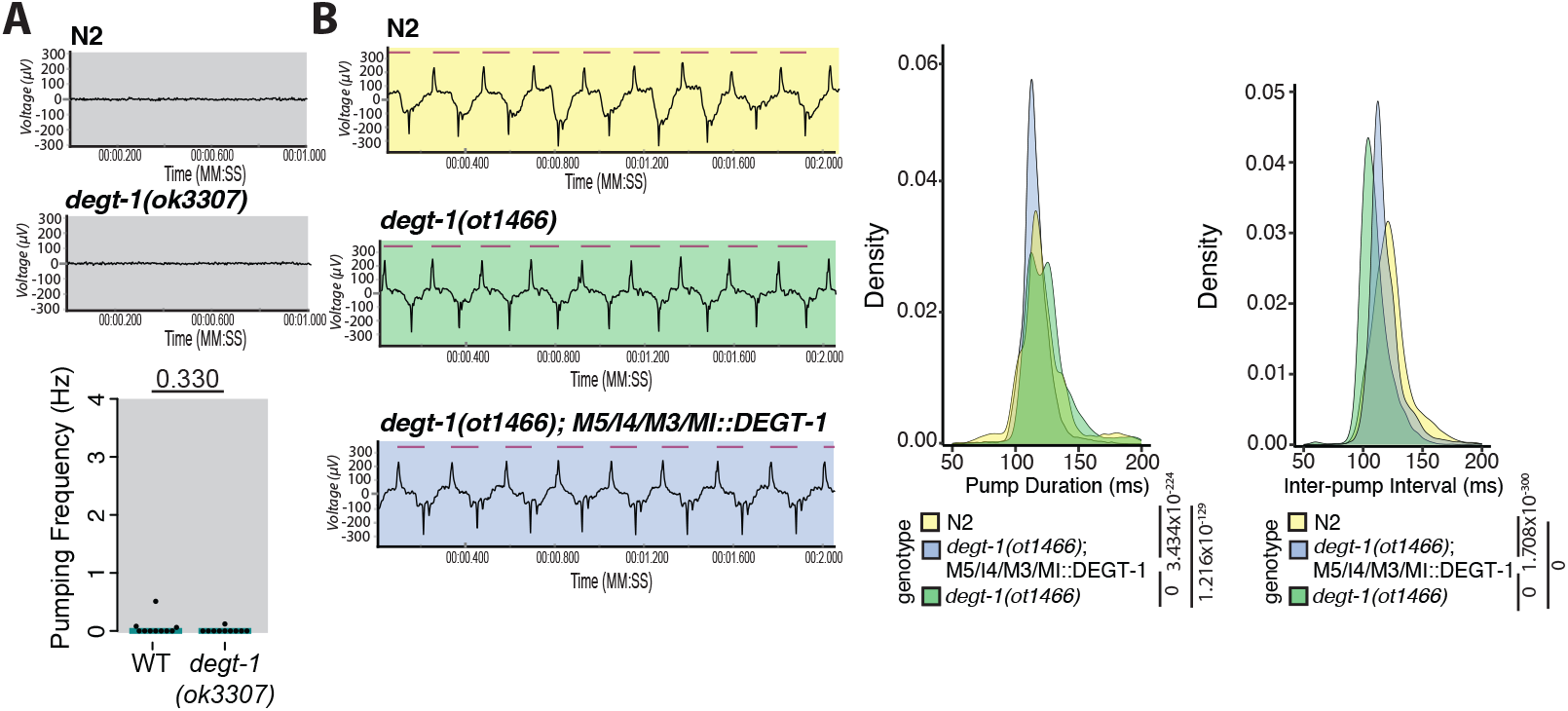
DEGT-1 mutants have a shortened inter-pump interval. **A:** *degt-1* mutants do not show promiscuous pumping in M9 (buffer control) in the ScreenChip. Sample electropharyngeogram traces for N2 and *degt-1(ok3307)* are shown above, quantification of pumping frequency per animal is shown below. Each black dot represents one animal, cyan bars show median. p-value (above plot) was calculated by Wilcoxon rank-sum test. **B:** *degt-1* mutants display altered pumping dynamics in response to 40mM serotonin in M9 in ScreenChip. Sample electropharyngeogram traces for N2, *degt-1(ot1466)*, and rescue are shown to the left. The pink bars above the trace represent the pump duration (as annotated by the NemAnalysis software), the length between each pink bar is each inter-pump interval. Density plots of pump duration and inter-pump interval are shown to the right. Rescue line 1 (**Fig. 3D**) was chosen as representative for the ScreenChip assay, as it showed less phenotypic variability in the freely-feeding assay. N2 n=16,328 pumps from 22 animals. *degt-1(ot1466);* M5/I4/M3/MI::DEGT-1 n=16,434 pumps from 22 animals. *degt-1(ot1466)* n=15,449 pumps from 22 animals. X-axis display limits were set for both pump duration and inter-pump interval to visualize the differences between genotypes; the same data are shown with the complete range of x values in **Supp. Fig. 4B** (all statistical analysis was performed on the complete dataset). p-values (shown to the right of each genotype legend) were calculated using the Kolmogorov-Smirnov test.

Using 40mM serotonin as a pumping stimulus, we observed that wild-type animals display predominantly ∼115ms pumps, with occasional longer pumps (140ms; **Fig. 4B, Supp. Fig. 4A**). This is consistent with previous reports measuring pharyngeal pumping using the ScreenChip device, which showed that wild-type animals show a distribution of both shorter and longer pharyngeal pumps^36^. We found that in the absence of *degt-1* this distribution was altered; animals still displayed short and long pumps, but the duration of both of these pumps was slightly shorter (∼110ms and ∼130ms), and pumps were more equally distributed between the two durations, giving similar average pump durations of 122ms (N2) and 128ms (*degt-1* mutant). In our M3/MI/I4/M5::DEGT-1 rescuing line, which we had found fully rescued pharyngeal pumping frequency on food, the pump duration distribution is biased toward short pumps, as in wild-type animals.

The interval between pharyngeal pumps (inter-pump interval) showed a unimodal distribution, also consistent with previous reports^36^. In wild-type animals this distribution was centered around ∼125ms, but in *degt-1* mutants the inter-pump interval was significantly shorter (∼105ms; **Fig 4B, Supp. Fig. 4A**). In regards to inter-pump interval, our M3/MI/I4/M5::DEGT-1 rescuing line also showed an intermediate value relative to wild-type and the *degt-1* mutant (∼115ms). While the alterations in pump duration between wild-type and *degt-1* mutants are more complex (shortening of duration of both shorter and longer pumps, but also a change in distribution between the two pump lengths), the change in inter-pump interval in a *degt-1* mutant represents a 20% shortening. In the context of our results when animals were freely feeding (wild-type at ∼4.75-5Hz, *degt-1* mutants at ∼5.75-6.25Hz), a 20% shortening would accomplish an increase of ∼1Hz. We thus conclude that the overall faster feeding rate that we observed in animals freely moving on food is predominantly the result of a DEGT-1 function in the I4 and M5 neurons to regulate the duration of the inter-pump interval.

Based on the electron micrograph reconstruction of the pharyngeal nervous system, neither the I4 nor M5 neurons have direct contact with the pharyngeal lumen^12^. Additionally, the distinct colocalization of the DEGT-1 protein with the pharyngeal basement membrane suggested to us that the I4 and M5 neurons are responding to some type of mechanosensory stimulus other than ingested food (or distension of the pharyngeal lumen) to shape pharyngeal pumping. We wondered whether previously unappreciated mechanosensory forces occur during a pharyngeal pump that could be sensed by the I4 and M5 neurons. High-speed video has been generated of *C. elegans* pharyngeal pumping at 1000 frames-per-second (fps)^38^, and we noticed in these videos that the posterior bulb of the pharynx undergoes significant anterior-posterior movement within the head of the animal during a pharyngeal pump. We analyzed this movement by quantifying anterior-posterior movement of the posterior bulb relative to the rest of the head. This analysis demonstrated that the pharynx indeed moves within the head of the animal during the course of pharyngeal pumping (**Supplemental Figure 4B**). Specifically, we noted that the posterior bulb shifts posteriorly during the grinder inversion step of pumping. Grinder inversion is accomplished by contraction of the posterior bulb muscles, which flips and compresses the chitinous grinder plates and serves to break open the ingested bacterial cells^13^. We found that this contraction not only distorts the posterior bulb muscles at their luminal surfaces, but also at the surface apposing the pharyngeal basement membrane; this surface is where the I4 and M5 neuronal soma are located, and where we observed DEGT-1 protein localization (**Fig. 1E**). The location of DEGT-1 protein in the I4 and M5 neurons with respect to the pharyngeal basement membrane suggest that muscle movements during the grinding step of pharyngeal pumping create shear forces at the border of the pharynx. We propose that these are sensed via DEGT-1 in the I4 and M5 neurons to provide feedback of a completed pharyngeal pumping cycle. In the absence of this feedback, the organ overcompensates for the perceived lack of pharyngeal pumping by increasing the pumping rate.

## DISCUSSION

Our study shows that the DEG/ENaC channel DEGT-1 is expressed exclusively in 4 pharyngeal neuron classes, localizes to the pharyngeal basement membrane, and is required in the I4 and M5 neurons to regulate the frequency of pharyngeal pumping. While mechanosensory neurons are known to be required for ingestion and digestion across many animals, it has been much more difficult to identify the molecular machinery governing enteric mechanosensation^5^.

In cases where DEG/ENaC channels have been implicated in enteric mechanosensation, the relevant channels sense either the ingested contents per se^7^, or distension caused by the bolus^8,9^. Based on four independent lines of evidence we propose that DEGT-1, in contrast, functions as a proprioceptive channel in I4 and M5 to regulate ingestion. First, DEGT-1 protein localizes to the neuronal soma apposing the pharyngeal basement membrane, rather than neurites apposing the pharyngeal lumen. Second, we found that DEGT-1 was specifically required in the I4 and M5 neurons, neither of which have any anatomical contact to the pharyngeal lumen or ingested food. Third, *degt-1* knockout results in alterations to pharyngeal pumping both when animals are assessed freely feeding, or when using serotonin as the stimulus (thus, not ingesting any food). Fourth, we found that the grinding step of each pharyngeal pump, previously proposed to be controlled by the M5 neuron, displaces the posterior bulb relatively to the head. This muscle movement provides a source of shear force at the border of the pharynx that could be detected by DEGT-1 in both of the DEGT-1+ posterior bulb neurons.

Another recently-described aspect of pharyngeal pumping, the ‘pharyngeal plunge,’ occurs during a subset of pharyngeal pumps and is regulated by the pharyngeal-intestinal valve(vpi)^39^. This movement involves the posterior bulb depressing into the anterior intestine. We believe the posterior bulb movements during grinding that we observed to be distinct from the pharyngeal plunge, which is reported to occur only once every ∼4 seconds (or 16-20 pumps), and be a large displacement (5 microns) of the pharynx^39^. Movement of the grinder during a pharyngeal pump is a well-described phenomenon, and can be readily quantified both automatically and by eye^35^, but the resulting pharyngeal displacement had not been described.

In the case of the M5 neuron, our data provide experimental validation for the grinding function that had been proposed computationally based on network analysis^12^. Specifically, M5 is the only neuron that innervates the posterior-most pharyngeal muscles, which control the contraction that leads to grinder inversion. Based on the localization of DEGT-1, we propose that M5 responds to the force generated by this posterior displacement of the pharynx, and this information is used by the animal to determine that a pump has been successfully performed. The I4 neuron, which we also found to be necessary and sufficient for regulating pharyngeal pumping frequency via DEGT-1, has been proposed to serve a neuromodulatory role^12^. I4 may respond to proprioceptive input by releasing a neuropeptide that modulates either the function of other pharyngeal neurons, or the pharyngeal muscle directly. Because restoring DEGT-1 to I4 alone fully rescues the rapid pumping phenotype, we do not expect that I4 simply acts upstream of M5 in a linear circuit. Consistent with this, I4 does not have direct chemical synaptic connectivity with M5^10,12^. Thus, I4 and M5 may be acting fully redundantly to regulate pumping frequency via DEGT-1 (ie neuronal degeneracy), a feature which has been proposed to be widespread in the pharyngeal nervous system to ensure robustness of function^35^.

While DEGT-1 is also expressed in and has a similar protein localization pattern in the MI and M3 neurons, we did not find that its expression in either of these neurons was sufficient or necessary for normal pumping. Like I4 and M5, the MI neuron does not have a known function^12^, while M3 has been shown to act as a proprioceptor for pharyngeal muscle contraction^14^, and regulate the duration of pharyngeal pumping via cholinergic neurotransmission^29^. However, in contrast to the localization of DEGT-1 protein at the pharyngeal basement membrane, the M3 neuron has also been shown to have putative sensory endings exposed to the pharyngeal lumen itself^12^. This suggests that the M3 ablation phenotype may be the result of a function in direct detection of ingested food, or sensation of the resulting change in width of the pharyngeal lumen itself. The expression of DEGT-1 in MI and M3 may still suggest that these neurons respond to mechanosensory forces, even those generated at the pharyngeal basement membrane. However, it is possible that the purpose of this mechanosensation is for some role other than regulating pumping frequency, or that it is important only in some specific environmental or behavioral context.

We found that local proprioception in the *C. elegans* foregut not only feeds back onto feeding rate, but also serves as an important checkpoint to regulate animal size and lipid storage. In the absence of *degt-1*, excess feeding results in adult animals accumulating increased lipid and consequently, premature senescence phenotypes. This type of homeostatic overcompensation has been observed in multiple contexts when a loss of mechanosensation abrogates proprioceptive feedback, such as excessive body bending during locomotion in *C. elegans* lacking the *trp-4* TRPN channel^40^, or hypermetric movements in human patients lacking PIEZO2^41^. Animals use multiple mechanical and chemosensory cues to judge the quantity and quality of meals, such as nutrient content and ingested volume (via distension)^7,42,43^. We show here that it is also crucial for the feeding organ to sense its own ingestive movements, and that dysregulation of this proprioception has global effects on animal health and aging.

## Acknowledgements

We thank Chi Chen for generating transgenic strains, and Wendy Cao for providing the *otEx8085[pha-4prom2::GFP, pha-1 rescuing DNA]* reporter, and for communicating unpublished results on promoters specific for each pharyngeal neuron. Strains were also provided by the CGC, which is supported by the NIH Office of Research Infrastructure Programs (P40 OD010440). This research was supported by the Jane Coffin Childs Memorial Fund for Medical Research (EAB) and the University of Basel (AFS).

## REFERENCE

1. Mazzuoli, G. & Schemann, M. Mechanosensitive enteric neurons in the myenteric plexus of the mouse intestine. PLoS One 7, e39887 (2012). 10.1371/journal.pone.0039887

2. Kugler, E. M. et al. Mechanical stress activates neurites and somata of myenteric neurons. Front Cell Neurosci 9, 342 (2015). 10.3389/fncel.2015.00342

3. Mazzuoli-Weber, G. et al. Piezo proteins: incidence and abundance in the enteric nervous system. Is there a link with mechanosensitivity? Cell Tissue Res 375, 605–618 (2019). 10.1007/s00441-018-2926-7

4. Gottlieb, P. et al. Revisiting TRPC1 and TRPC6 mechanosensitivity. Pflugers Arch 455, 1097–1103 (2008). 10.1007/s00424-007-0359-3

5. Mercado-Perez, A. & Beyder, A. Gut feelings: mechanosensing in the gastrointestinal tract. Nat Rev Gastroenterol Hepatol 19, 283–296 (2022). 10.1038/s41575-021-00561-y

6. Bianchi, L. & Driscoll, M. Protons at the gate: DEG/ENaC ion channels help us feel and remember. Neuron 34, 337–340 (2002). 10.1016/s0896-6273(02)00687-6

7. Rhoades, J. L. et al. ASICs Mediate Food Responses in an Enteric Serotonergic Neuron that Controls Foraging Behaviors. Cell 176, 85–97 e14 (2019). 10.1016/j.cell.2018.11.023

8. Olds, W. H. & Xu, T. Regulation of food intake by mechanosensory ion channels in enteric neurons. Elife 3 (2014). 10.7554/eLife.04402

9. Page, A. J. et al. Different contributions of ASIC channels 1a, 2, and 3 in gastrointestinal mechanosensory function. Gut 54, 1408–1415 (2005). 10.1136/gut.2005.071084

10. Albertson, D. G. & Thomson, J. N. The pharynx of Caenorhabditis elegans. Philos Trans R Soc Lond B Biol Sci 275, 299–325 (1976). 10.1098/rstb.1976.0085

11. Keeley, D. P. et al. Comprehensive Endogenous Tagging of Basement Membrane Components Reveals Dynamic Movement within the Matrix Scaffolding. Dev Cell 54, 60–74 e67 (2020). 10.1016/j.devcel.2020.05.022

12. Cook, S. J. et al. The connectome of the Caenorhabditis elegans pharynx. J Comp Neurol 528, 2767–2784 (2020). 10.1002/cne.24932

13. Avery, L. & Horvitz, H. R. Pharyngeal pumping continues after laser killing of the pharyngeal nervous system of C. elegans. Neuron 3, 473–485 (1989). 10.1016/0896-6273(89)90206-7

14. Avery, L. Motor neuron M3 controls pharyngeal muscle relaxation timing in Caenorhabditis elegans. J Exp Biol 175, 283–297 (1993). 10.1242/jeb.175.1.283

15. Iwanir, S. et al. Serotonin promotes exploitation in complex environments by accelerating decision-making. BMC Biol 14, 9 (2016). 10.1186/s12915-016-0232-y

16. Taylor, S. R. et al. Molecular topography of an entire nervous system. Cell 184, 4329–4347 e4323 (2021). 10.1016/j.cell.2021.06.023

17. O’Hagan, R., Chalfie, M. & Goodman, M. B. The MEC-4 DEG/ENaC channel of Caenorhabditis elegans touch receptor neurons transduces mechanical signals. Nat Neurosci 8, 43–50 (2005). 10.1038/nn1362

18. Huang, M. & Chalfie, M. Gene interactions affecting mechanosensory transduction in Caenorhabditis elegans. Nature 367, 467–470 (1994). 10.1038/367467a0

19. Shi, S., Luke, C. J., Miedel, M. T., Silverman, G. A. & Kleyman, T. R. Activation of the Caenorhabditis elegans Degenerin Channel by Shear Stress Requires the MEC-10 Subunit. J Biol Chem 291, 14012–14022 (2016). 10.1074/jbc.M116.718031

20. Chelur, D. S. et al. The mechanosensory protein MEC-6 is a subunit of the C. elegans touch-cell degenerin channel. Nature 420, 669–673 (2002). 10.1038/nature01205

21. Liu, J., Schrank, B. & Waterston, R. H. Interaction between a putative mechanosensory membrane channel and a collagen. Science 273, 361–364 (1996). 10.1126/science.273.5273.361

22. Jospin, M. & Allard, B. An amiloride-sensitive H+-gated Na+ channel in Caenorhabditis elegans body wall muscle cells. J Physiol 559, 715–720 (2004). 10.1113/jphysiol.2004.069971

23. Du, H., Gu, G., William, C. M. & Chalfie, M. Extracellular proteins needed for C. elegans mechanosensation. Neuron 16, 183–194 (1996). 10.1016/s0896-6273(00)80035-5

24. Graham, P. L. et al. Type IV collagen is detectable in most, but not all, basement membranes of Caenorhabditis elegans and assembles on tissues that do not express it. J Cell Biol 137, 1171–1183 (1997). 10.1083/jcb.137.5.1171

25. Jayadev, R. et al. alpha-Integrins dictate distinct modes of type IV collagen recruitment to basement membranes. J Cell Biol 218, 3098–3116 (2019). 10.1083/jcb.201903124

26. Sibley, M. H., Graham, P. L., von Mende, N. & Kramer, J. M. Mutations in the alpha 2(IV) basement membrane collagen gene of Caenorhabditis elegans produce phenotypes of differing severities. EMBO J 13, 3278–3285 (1994). 10.1002/j.1460-2075.1994.tb06629.x

27. Driscoll, M. & Chalfie, M. The mec-4 gene is a member of a family of Caenorhabditis elegans genes that can mutate to induce neuronal degeneration. Nature 349, 588–593 (1991). 10.1038/349588a0

28. Tavernarakis, N. & Driscoll, M. Molecular modeling of mechanotransduction in the nematode Caenorhabditis elegans. Annu Rev Physiol 59, 659–689 (1997). 10.1146/annurev.physiol.59.1.659

29. Dent, J. A., Davis, M. W. & Avery, L. avr-15 encodes a chloride channel subunit that mediates inhibitory glutamatergic neurotransmission and ivermectin sensitivity in Caenorhabditis elegans. EMBO J 16, 5867–5879 (1997). 10.1093/emboj/16.19.5867

30. O’Rourke, E. J., Soukas, A. A., Carr, C. E. & Ruvkun, G. C. elegans major fats are stored in vesicles distinct from lysosome-related organelles. Cell Metab 10, 430–435 (2009). 10.1016/j.cmet.2009.10.002

31. Zhao, Y. et al. Two forms of death in ageing Caenorhabditis elegans. Nat Commun 8, 15458 (2017). 10.1038/ncomms15458

32. Van Gilst, M. R., Hadjivassiliou, H., Jolly, A. & Yamamoto, K. R. Nuclear hormone receptor NHR-49 controls fat consumption and fatty acid composition in C. elegans. PLoS Biol 3, e53 (2005). 10.1371/journal.pbio.0030053

33. Herndon, L. A. et al. Stochastic and genetic factors influence tissue-specific decline in ageing C. elegans. Nature 419, 808–814 (2002). 10.1038/nature01135

34. Raizen, D. M., Lee, R. Y. & Avery, L. Interacting genes required for pharyngeal excitation by motor neuron MC in Caenorhabditis elegans. Genetics 141, 1365–1382 (1995). 10.1093/genetics/141.4.1365

35. Trojanowski, N. F., Padovan-Merhar, O., Raizen, D. M. & Fang-Yen, C. Neural and genetic degeneracy underlies Caenorhabditis elegans feeding behavior. J Neurophysiol 112, 951–961 (2014). 10.1152/jn.00150.2014

36. Millet, J. R. M., Romero, L. O., Lee, J., Bell, B. & Vasquez, V. C. elegans PEZO-1 is a mechanosensitive ion channel involved in food sensation. J Gen Physiol 154 (2022). 10.1085/jgp.202112960

37. Raizen, D. M. & Avery, L. Electrical activity and behavior in the pharynx of Caenorhabditis elegans. Neuron 12, 483–495 (1994). 10.1016/0896-6273(94)90207-0

38. Brenner, I. R., Raizen, D. M. & Fang-Yen, C. Pharyngeal timing and particle transport defects in Caenorhabditis elegans feeding mutants. J Neurophysiol 128, 302–309 (2022). 10.1152/jn.00444.2021

39. Park, Y. J. et al. PIEZO acts in an intestinal valve to regulate swallowing in C. elegans. Nat Commun 15, 10072 (2024). 10.1038/s41467-024-54362-3

40. Li, W., Feng, Z., Sternberg, P. W. & Xu, X. Z. A C. elegans stretch receptor neuron revealed by a mechanosensitive TRP channel homologue. Nature 440, 684–687 (2006). 10.1038/nature04538

41. Chesler, A. T. et al. The Role of PIEZO2 in Human Mechanosensation. N Engl J Med 375, 1355–1364 (2016). 10.1056/NEJMoa1602812

42. Dockray, G. J. Enteroendocrine cell signalling via the vagus nerve. Curr Opin Pharmacol 13, 954–958 (2013). 10.1016/j.coph.2013.09.007

43. Bai, L. et al. Genetic Identification of Vagal Sensory Neurons That Control Feeding. Cell 179, 1129–1143 e1123 (2019). 10.1016/j.cell.2019.10.031

44. Hobert, O. The neuronal genome of Caenorhabditis elegans. WormBook, 1–106 (2013). 10.1895/wormbook.1.161.1

45. Jin, E. J. & Jin, Y. A mutation linked to degt-1(ok3307) in C. elegans strain VC2633 affects rpm-1. MicroPubl Biol 2022 (2022). 10.17912/micropub.biology.000565

