## Supplemental Figures and Legends, Methods for "The mechanosensory DEG/ENaC channel DEGT-1 is a proprioceptor of *C. elegans* foregut movement"

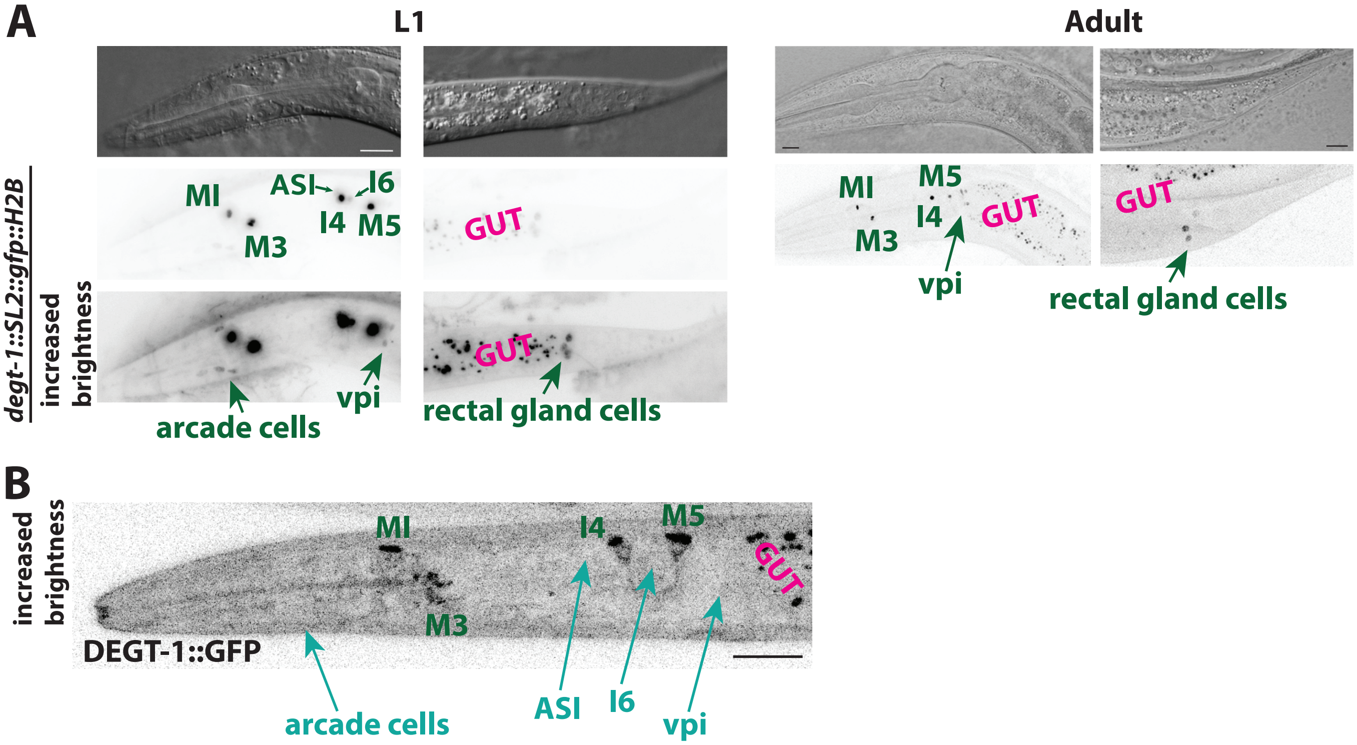


Supplemental Figure 1

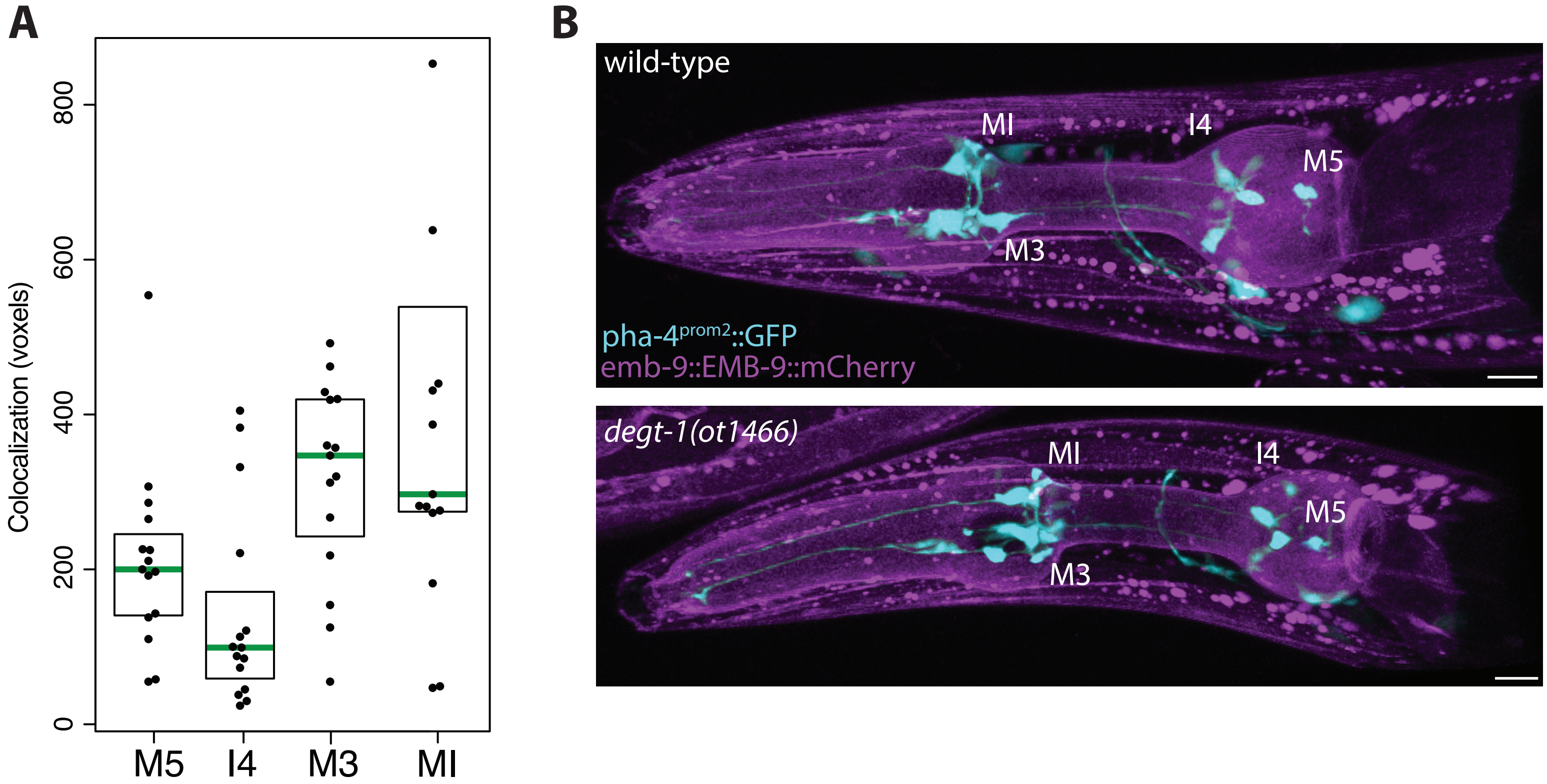


Supplemental Figure 2

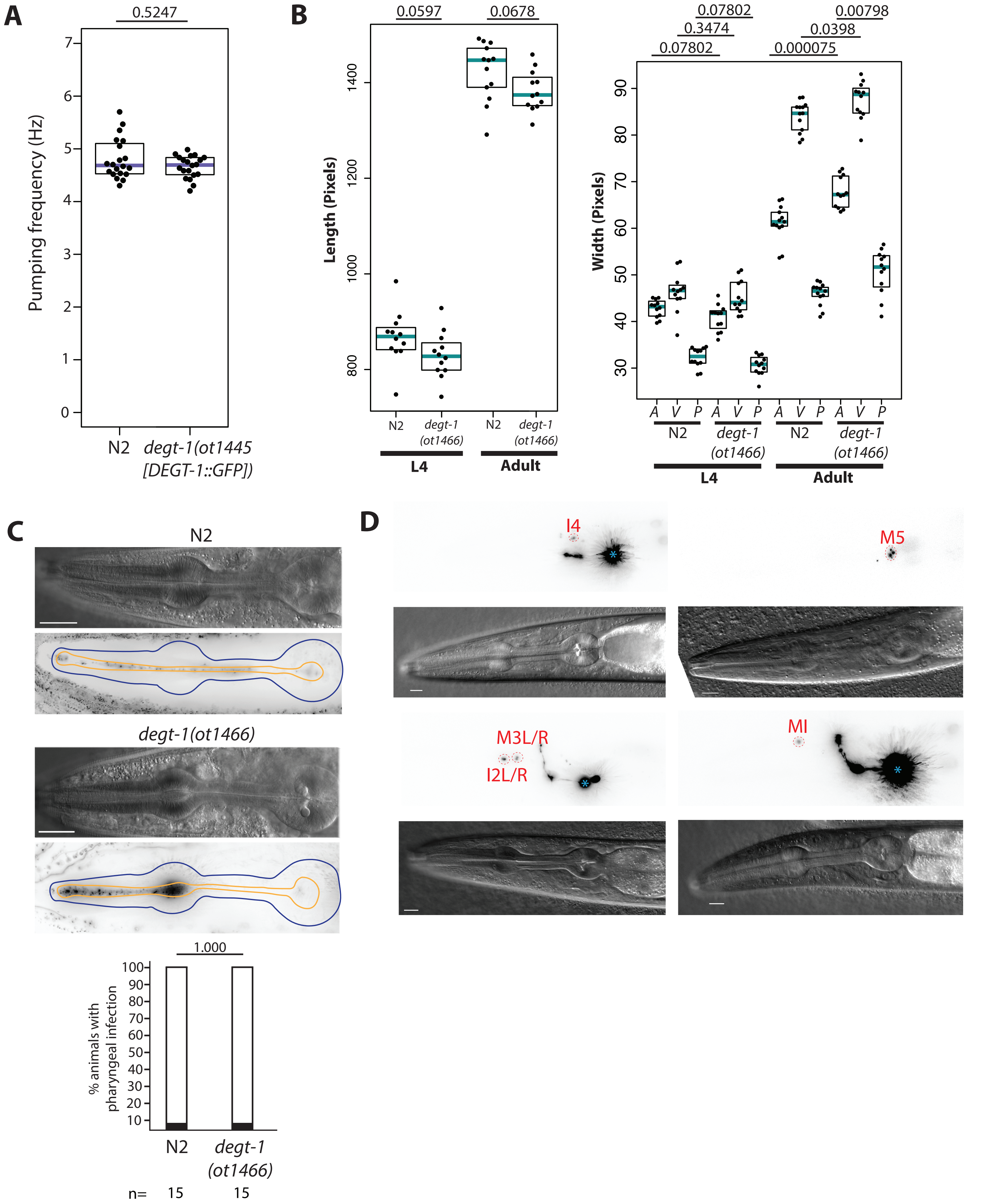


Supplemental Figure 3

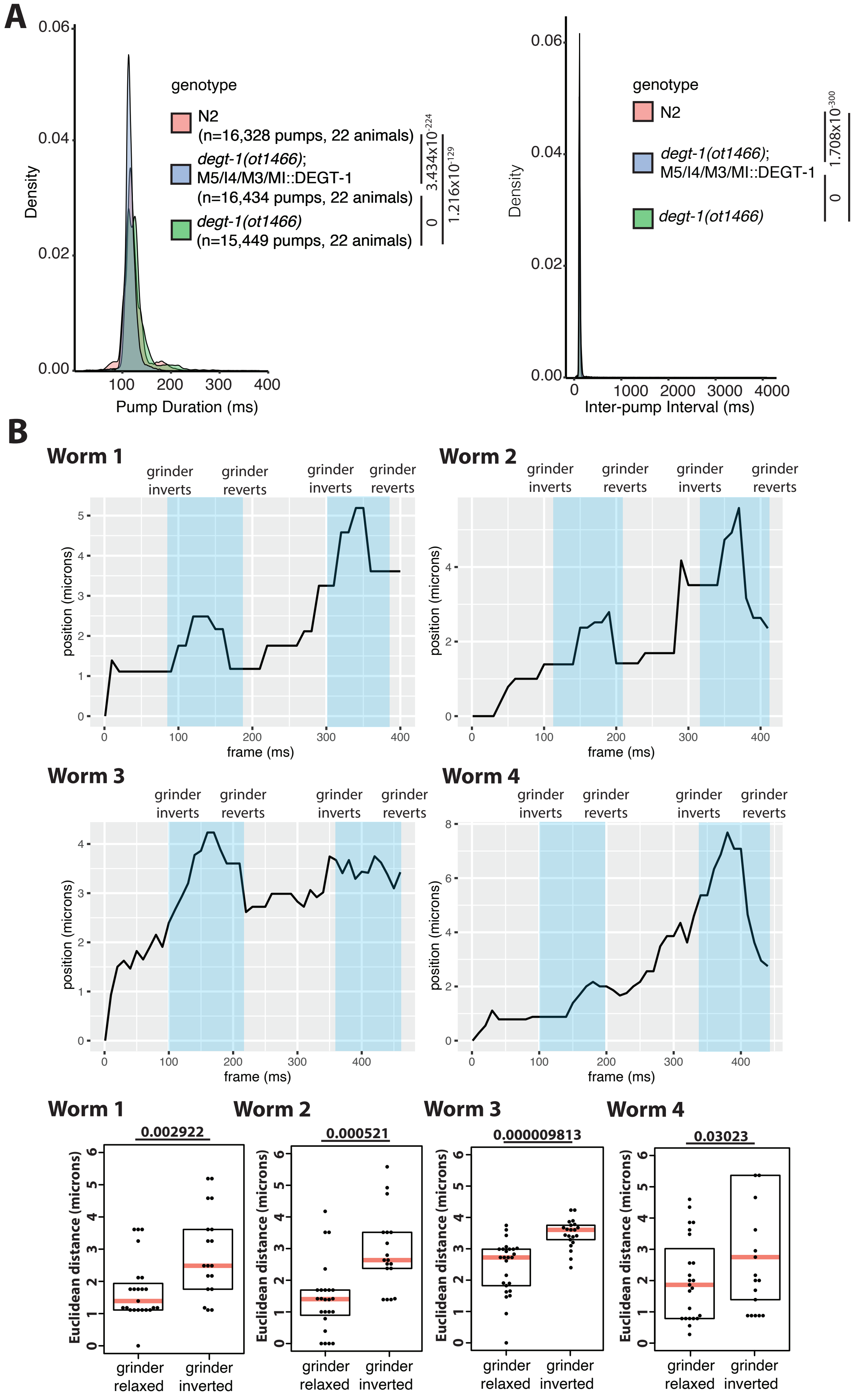


Supplemental Figure 4

### SUPPLEMENTAL FIGURE LEGENDS

**Supplemental Figure 1 (related to Figure 1): A *degt-1* transcriptional reporter shows strong expression in MI/M3/I4/M5, and additional weak/variable expression.**

**A:** Expression of *degt-1::SL2::gfp::H2B (syb9045)* transcriptional reporter in L1 and adult animals. DIC images shown above, GFP expression (black and white inverted) shown below. Compared to the translational reporter, we observed some additional expression in this reporter. In every animal scored in L1, L4, and adult stages, we could observe expression in the pharyngeal-intestinal valve (vpi) cells, and in the three rectal gland cells. In L1 animals, we observed consistent expression in the anterior-most pair of intestinal cells, but this vanished across developmental time. Stringent CeNGEN thresholds (3 and 4) predict expression exclusively in MI, M3, I4, and M5. We sometimes observed extremely dim expression in neurons consistent with the lower two CeNGEN thresholds (1 and 2): I6 and ASI, and in a variable number of cells consistent with the arcade cells. We never saw expression in I3, an additional neuron reported by these thresholds. We were never able to recapitulate this additional dim/variable/inconsistent transcriptional expression with our translational reporter (ie **Supplemental Fig 1B,** maximum intensity projection). This could suggest that *degt-1* is transcribed (and spliced) but not translated in these cells, or subjected to post-transcriptional or post-translational degradation. For the L1 animal, the same images are shown both at normal brightness (to show nervous system expression) and at increased brightness (to show much dimmer non-neuronal expression). While previous studies using promoter fusion reporters have suggested that *degt-1* is expressed more broadly, including in the PVD neuron[1,2], we did not observe PVD expression in either of our endogenously-tagged reporters at any developmental stage, nor is it reported in single cell RNA sequencing data[3].

**B:** Expression of DEGT-1::GFP (*ot1445*) translational reporter (black and white inverted) in an L1 animal. Same image shown as **Fig. 1E,** brightness increased to emphasize the lack of additionally expressing cells. The four expressing pharyngeal neurons are labeled in green, while the cells indicated with teal arrows show transcriptional but not translational expression of *degt-1*. All scale bars 10uM. Autofluorescent gut granules are labeled GUT (magenta) in both **A** and **B**.

**Supplement 2 (related to Figure 2): DEGT-1 colocalizes with the pharyngeal basement membrane, but is not required for neuronal attachment.**

**A:** Quantification of DEGT-1::GFP colocalization with EMB-9::mCherry. Colocalization was quantified in voxels using Imaris Analysis (see Methods). Each black dot represents colocalization region from one animal, green bar indicates median, black boxes indicate quartiles.

**B:** DEGT-1 is not required for anchoring neurons to the pharyngeal basement membrane. The soma of the four DEGT-1+ neurons were assessed for colocalization with the pharyngeal basement membrane using a cytoplasmic GFP marker in wild-type and *degt-1* mutant animals. 15 adult animals were analyzed for each genotype, and colocalization was observed in all *degt-1*-expressing neurons in 100% of animals in both genotypes. Representative volumetric projections are shown of both wild-type and mutant animals, with *degt-1*-expressing neurons labeled. Scale bars (white) are 10uM.

**Supplemental Figure 3 (related to Figure 3): Loss of DEGT-1 affects animal width but not pharyngeal health, and can be cell-specifically rescued.**

**A:** The DEGT-1::GFP fusion protein does not abrogate *degt-1* function. Each black dot is the pharyngeal pumping frequency of one animal, purple bars are median, black boxes are quartiles. p-value (above plot) calculated using Wilcoxon rank-sum test.

**B:** *degt-1* mutants are wider, but not longer, than wild-type animals as day 1 adults, but not L4s. Each black dot represents the measured length or width of one animal, cyan bars are medians, black boxes are quartiles. p-values (above plot) calculated using Wilcoxon rank-sum test. For width plot (right), width was measured at 3 points on each animal: A=anterior, V=vulva, P=posterior (see Methods).

**C:** *degt-1* mutants do not display enhanced pharyngeal infection at day 5 of adulthood. Representative DIC images (above) and black and white inverted GFP (below) of wild-type and *degt-1(ot1466)* Fluorescent bacteria are visible within the pharyngeal lumen (yellow border) but have not invaded the pharyngeal muscle (between the yellow and blue borders). Scale bars (white) are 20uM. p-value shown above the plot was calculated with Fisher's Exact Test.

**D:** Expression of rescuing constructs in day 1 adult animals (black and white inverted mCherry::H2B), with DIC channel shown below. Neurons are indicated with red dashed circles. The *aqp-5* promoter is expressed in the normally DEGT-1- neuron I2 in addition to the DEGT-1+ neuron M3, but this additional expression did not confer any pumping phenotype in feeding animals (**Fig. 3D**). Scale bars are 10uM. The ttx-3::mCherry transgene was used as a coinjection marker, so the AIY interneuron is also visible in some images (marked with a blue asterisk).

**Supplement 4 (related to Figure 4): Displacement of posterior bulb during grinder inversion suggests sensory modality for *degt-1* mutant phenotype.**

**A:** *degt-1* mutants have shortened inter-pump intervals. The same ScreenChip data from **Fig. 4B**, with the complete range of x values shown. While duration of a pharyngeal pump exhibits a fairly narrow distribution, inter-pump interval takes into account not only intervals between pumps when animals are feeding rapidly, but also pauses in feeding during the recording are reported as a prolonged inter-pump interval. Thus, the plot is stretched by occasional inter-pump intervals as long as 4000ms, although the vast majority of the data is represented in the region shown in **Fig. 4B** (50-200ms)**.**

**B:** The posterior bulb is displaced within the head of the animal during grinder inversion. High-speed recordings of pumping from four wild-type animals[4] were analyzed (each containing two pharyngeal pumps; see Methods). Above, a trace of the Euclidean distance between the centroid of the posterior bulb and an anchor point on the head of the animal across the length of the entire recording is shown for each animal. Above each trace, the points in each video when the grinder inverts and then reverts are labeled, and the time that the grinder is inverted is indicated in the blue box. Below, the Euclidean distance between the posterior bulb and head anchor point at frames when the grinder is inverted versus frames when the grinder is relaxed for the entire video are plotted for each animal. Each black dot represents one Euclidean distance, salmon bars are median, black boxes are quartiles. p-values (above plot) were calculated using the Wilcoxon rank-sum test.

### METHODS

#### Strains

Worms were maintained by standard methods[5]. Worms were grown at 20°C on nematode growth media (NGM) plates seeded with bacteria (*E.coli* OP50) as a food source, except where explicitly indicated otherwise. The following alleles and transgenes were used in this study:

| Strain Name | Alleles/Transgenes | Expression/notes | Reference | Figures |
|---|---|---|---|---|
| OH18902 | *degt-1(ot1445)* | Translational DEGT-1::GFP | This study | 1D,E, Supp. Fig. 1B, Supp. Fig. 3A |
| PHX9045 | *degt-1(syb9045)* | Transcriptional *degt-1::SL2::GFP::H2B* | This study | 1D, Supp. Fig. 1A |
| OH19034 | *degt-1(ot1466)* | CRISPR-generated null | This study | 1D, 3A, 3B, 3C, 3D, Supp. Fig. 3B, Supp. Fig. 3C, 4B, Supp. Fig. 4A |
| CZ29001 | *degt-1(ok3307)* | Deletion mutant | [6] | 1D, 4A |

| | | | | |
|---|---|---|---|---|
| OH19286 | *qyIs46[emb-9p::emb-9::mCherry + unc-119(+)]; degt-1(ot1445)* | Basement membrane collagen marker; DEGT-1::GFP | [7] and this study | 2A, 2B, Supp. Fig. 2A |
| OH19315 | *let-2(ot1542); degt-1(ot1445); qyIs45* | *ot1542* is a CRISPR-generated temperature-sensitive allele identical to *let-2(g30)* | [8] | 2B |
| OH19551 | *qyIs46; otEx8085[pha-4prom2::GFP, pha-1 rescuing DNA]* | *pha-4* promoter with expression specific to all pharyngeal neurons and the RIS neuron | This study | Supp. Fig. 2B |
| OH19552 | *degt-1(ot1466); qyIs46; otEx8085* | | This study | Supp. Fig. 2B |
| N2 | | wild-type | [5] | 3A, 3B, 3C, 3D, Supp. Fig. 3A, Supp Fig. 3B, Supp. Fig. 3C, 4A, 4B, Supp. Fig. 4A, Supp. Fig. 4B |

| | | | | |
|---|---|---|---|---|
| OH19285 | *degt-1(ot1466); otEx8251[sams-5::degt-1::SL2::mCherry, clec-166::degt-1::SL2::mCherry, aqp-5::degt-1::SL2::mCherry, pha-2GFPF::degt-1::SL2::mCherry, ttx-3::mCherry]* | M3/MI/I4/M5 rescue line 1 | This study | 3D, 4B, Supp. Fig. 4A |
| OH19284 | *degt-1(ot1466); otEx8250[sams-5::degt-1::SL2::mCherry, clec-166::degt-1::SL2::mCherry, aqp-5::degt-1::SL2::mCherry, pha-2GFPF::degt-1::SL2::mCherry, ttx-3::mCherry]* | M3/MI/I4/M5 rescue line 2 | This study | 3D |
| OH19293 | *degt-1(ot1466); otEx8252[pha-2GFP-F::degt-1::SL2::mCherry, ttx-3::mCherry]* | I4 rescue line 1 | This study | 3D, Supp. Fig. 3D |
| OH19294 | *degt-1(ot1466); otEx8253[pha-2GFP-F::degt-* | I4 rescue line 2 | This study | 3D |

| | | | | |
|---|---|---|---|---|
| | *1::SL2::mCherry, ttx-3::mCherry]* | | | |
| OH19515 | *degt-1(ot1466); otEx8279[clec-166::degt-1::SL2::mCherry, ttx-3::mCherry]* | M5 rescue line 1 | This study | 3D |
| OH19301 | *degt-1(ot1466); otEx8254[clec-166::degt-1::SL2::mCherry, ttx-3::mCherry]* | M5 rescue line 2 | This study | 3D, Supp. Fig. 3D |
| OH19302 | *degt-1(ot1466); otEx8255[clec-166::degt-1::SL2::mCherry, ttx-3::mCherry]* | M5 rescue line 3 | This study | 3D |
| OH19306 | *degt-1(ot1466); otEx8256[sams-5::degt-1::SL2::mCherry, ttx-3::mCherry]* | MI rescue line 1 | This study | 3D, Supp. Fig. 3D |
| OH19307 | *degt-1(ot1466); otEx8257[sams-5::degt-1::SL2::mCherry, ttx-3::mCherry]* | MI rescue line 2 | This study | 3D |
| OH19311 | *degt-1(ot1466); otEx8258[aqp-5::degt-* | M3 rescue line 1 | This study | 3D, Supp. Fig. 3D |

| | | | | |
|---|---|---|---|---|
| | *1::SL2::mCherry, ttx-3::mCherry]* | | | |
| OH19312 | *degt-1(ot1466); otEx8259[aqp-5::degt-1::SL2::mCherry, ttx-3::mCherry]* | M3 rescue line 2 | This study | 3D |

**Cloning and constructs**

To generate *degt-1* rescuing constructs, the *degt-1* cDNA was synthesized (TWIST BIOSCIENCE) and assembled using NEBuilder HiFi Assembly Mix with an operon linker, mCherry::H2B (amplified from AddGene ID 89368), with SphI/XmaI restriction digest sites to insert the respective neuron-specific promoter fragments 5' of the *degt-1* cDNA. The following promoters were used to generate neuron-specific rescuing constructs:

The *aqp-5* promoter for M3 is 1777bp 5' of the *aqp-5* start codon.

The *sams-5* promoter for MI is 884bp 5' of the *sams-5* start codon.

The *clec-166* promoter for M5 is 765bp 5' of the *clec-166* start codon.

The *pha-2 GFP-F* promoter for I4 is a 2.7kb region from the *pha-2* 5' UTR[9].

Each of these promoters was amplified from N2 genomic DNA with overhangs to the *degt-1* cDNA construct, and inserted using NEBuilder HiFi Assembly Mix into the SphI/XmaI sites.

**CRISPR/Cas9-mediated mutations**

All gRNAs were ordered as crRNA from IDT, and then complexed with Cas9 protein (IDT) and tracrRNA (IDT). CRISPR/Cas9 genome editing was based on a previously-described protocol[10]. In this protocol, a co-injection marker is used to single F1 animals where injection yielded successful array formation, and edited F2 progeny were identified by single worm PCR screening.

OH18902 *degt-1(ot1445[DEGT-1::GFP])* was generated in an N2 background using the gRNA actattacattagataacaa and an asymmetric dsDNA repair template to insert GFP (amplified from pPD95.75) in-frame immediately before the *degt-1* stop codon with 119bp homology arms. A point mutation was introduced into the *degt-1* 3' UTR in the repair template to destroy the PAM site (relative to the stop codon, base +56 was mutated from G to A).

PHX9045 *degt-1(syb9045)* was generated in an N2 background by SunyBiotech to insert SL2::GFP::H2B at the C-terminal end of *degt-1*.

OH19034 *degt-1(ot1466)* was generated in an N2 background using the 5' gRNA TGTAATCGAGACACCGTCAA and 3' gRNA ACTATTACATTAGATAACAA, and is a deletion from -84 (relative to start codon) to +58 (relative to stop codon).

The temperature-sensitive *let-2* allele *g30* has previously been described[8]; we recreated this same point mutation (*ot1542*) using CRISPR with oligo-mediated repair due to genetic linkage with the desired reporter strain. The gRNA sequence was tctggtcagccaggataccc, the repair oligo was tggtactcctggcaaaccggcgtctcccttttctcTtgggtatcctggctgaccagacagtcctggaaga (capital T is the C to T point mutation).

**Microscopy and image analysis**

Worms were anesthetized using 100mM of sodium azide (NaN3) and mounted on 5% agar on glass slides. Worms were analyzed by Nomarski optics and fluorescence microscopy at 40X magnification (exceptions detailed below), using a Zeiss LSM 880 confocal laser-scanning microscope, Zeiss LSM 980 confocal laser-scanning microscope, or Zeiss AxioImager Z.2. Multidimensional data was reconstructed as maximum intensity projections using Zeiss Zen, Fiji, or Imaris (Oxford Instruments) software.

For Supp. Fig. 2A, colocalization was quantified using Imaris Analysis software (Oxford Instruments). First, the GFP channel was used to create a mask of DEGT-1 localization in each individual DEGT-1+ neuron. Then, the colocalization tool was used to output the size of the basement membrane region (EMB-9::mCherry) that colocalized with the DEGT-1::GFP+ region of each neuron.

For Supp. Fig. 3B, animals were imaged at 10X magnification, and Fiji software was used to trace the length and width of animals using line segments. For width measurements, each animal was measured at three points: a line transecting the most anterior intestinal nuclei, a line transecting the vulva, and a line transecting the most posterior intestinal nuclei. These three measurements are reported separately for each animal.

**Figure preparation and statistics**

Plots were generated in R using the beeswarm and ggplot2 packages. Statistical tests as indicated in the figure legends were performed in R. Figures were prepared using Adobe Photoshop and Adobe Illustrator.

**Pumping in freely-moving animals on bacteria**

Worms were staged as day 1 adults by picking L4 animals the day before the assay, and allowing to develop overnight into adults at room temperature (~22C) for equilibration before the assay. Adults were then singled to individual plates before the assay, and allowed to equilibrate for 15 minutes before pumping was recorded. Each animal was observed for one minute, and number of pumps (judged by grinder displacement) were recorded with a manual counter. All experiments were performed blinded to animal genotype, and with two independent replicates, and controls were always performed in parallel.

**Oil Red O staining and analysis**

Oil Red O staining was performed according to published protocols[11]. Images were acquired using an OMAX 18MP USB 3.0 C-Mount Microscope Camera, mounted on a standard binocular dissecting microscope (Nikon SMZ645), using ToupLite software. Exposure and gain were set manually to allow the use of identical imaging settings and thus comparison across two independent replicates on different days. 20-21 animals were imaged per genotype for each replicate. Images were analyzed for red intensity of intestinal staining based on a published method[12]. In short, the RGB image was inverted (so that intensity of staining is reported as brightness, rather than darkness), and the 3 color channels were separated. The blue channel image was used to create a threshold mask of the intestine, and this was set as a region of interest. The region of interest was then applied to the red channel image, and mean intensity of the red channel was recorded per animal. Worms were staged as day 1 adults by picking L4 animals the day before the assay, and allowing to develop overnight into adults at room temperature (~22C) for equilibration before the assay.

**Temperature-sensitive *let-2***

Animals with a reporters for basement membrane collagen (emb-9::EMB-9::mCherry) and the DEGT-1::GFP protein were shifted from the permissive temperature (15C) at the L1 stage and then grown for 72 hours at the restrictive temperature (25C) before imaging, at which point they had reached the L4 stage. Both wild-type and *let-2* mutant animals were subjected to the same temperature shift paradigm. Some *let-2* mutant animals died as a result of the *let-2* mutation over the course of 72 hours at 25C (judged by developmental arrest and failure to respond to touch), and these were excluded from imaging. Colocalization between GFP and mCherry was assessed for each of the four DEGT-1+ neurons individually as a binary state (if DEGT-1::GFP contacted the basement membrane mCherry in any Z-slice, this was scored as colocalization).

**Pharyngeal infection assay**

Animals were grown on standard OP50 until the L4 stage, and then transferred to OP50-GFP[13]. Animals were maintained on OP50-GFP until the fifth day of adulthood, and then imaged to assess whether fluorescent bacteria had invaded the bacterial lumen, as previously described[14], which was recorded as a binary characteristic (pharynx with or without bacterial infection).

**Premature senescence assay**

To evaluate premature vacuole accumulation, N2 and *degt-1(ot1466)* animals were picked as L4 animals and then grown to day 5 adults at 20C. The posterior half (centered around the gonad arm) of each animal was imaged at 40X, and then animals were characterized for the presence or absence of coelomic vacuoles as a binary characteristic.

**ScreenChip**

Worms were staged as day 1 adults by picking L4 animals the day before the assay, and allowing to develop overnight into adults at room temperature (~22C) for equilibration before the assay. Immediately before the assay, animals were size-selected by isolating animals in M9 solution that had passed through a 70 micron filter but been trapped by a 50 micron filter, to ensure comparable snugness within the microfluidic chip (which affects voltage measurements). ScreenChip40 size microfluidic chips were used for all experiments. For the M9 buffer control assay, animals were loaded directly into the ScreenChip device (InVivo Biosystems), and for serotonin-induced pumping, animals were pre-incubated in 40mM serotonin in M9 for 30 minutes in the dark before loading into the ScreenChip device, and this was performed in two independent replicates. Each recording was 3 minutes in duration, and recordings were analyzed using the default parameters of the NemAnalysis software (v0.2) to extract timestamps of pump starts and ends. For **Fig. 4A,** the built-in ‘export results’ feature was used to export pumping frequency. To perform the pump-level analysis in **Fig. 4B and Supp. Fig. 4A**, timestamps of pump starts and ends were exported for each individual recording, and then the data were extracted to calculate pump durations (pump ends-pump starts), and inter-pump intervals (pump starts-pump ends). Beeswarm and density plots were generated using the ggplot2 package in R.

**Posterior bulb displacement measurements**

Existing high speed (1000fps) data[4] were re-analyzed to assess posterior bulb movements during pharyngeal pumping. An ellipse was manually drawn around the posterior bulb of each animal, and the position of the centroid of this ellipse was recorded every 10 ms. In parallel, a line transecting the grinder was drawn at the first frame, and this line was kept in a consistent position based on hypodermal landmarks, also in 10ms intervals. For each of these 10ms intervals, the Euclidean distance between the ellipse centroid and the line position was calculated: sqrt((line x position-bulb x position)^2+(line y position-bulb y position)^2). These Euclidean distances were plotted to generate the traces in **Supp. Fig. 4B.** The timestamps of grinder inversions were manually noted in the videos and used to split the Euclidean distances into 'grinder relaxed' and 'grinder inverted' categories, as represented in the beeswarm plots below.

### REFERENCE


1 Chatzigeorgiou, M. *et al.* Specific roles for DEG/ENaC and TRP channels in touch and thermosensation in C. elegans nociceptors. *Nat Neurosci* **13**, 861-868 (2010). https://doi.org/10.1038/nn.2581

2 Tao, L. *et al.* Parallel Processing of Two Mechanosensory Modalities by a Single Neuron in C. elegans. *Dev Cell* **51**, 617-631 e613 (2019). https://doi.org/10.1016/j.devcel.2019.10.008

3 Taylor, S. R. *et al.* Molecular topography of an entire nervous system. *Cell* **184**, 4329-4347 e4323 (2021). https://doi.org/10.1016/j.cell.2021.06.023

4 Brenner, I. R., Raizen, D. M. & Fang-Yen, C. Pharyngeal timing and particle transport defects in Caenorhabditis elegans feeding mutants. *J Neurophysiol* **128**, 302-309 (2022). https://doi.org/10.1152/jn.00444.2021

5 Brenner, S. The genetics of Caenorhabditis elegans. *Genetics* **77**, 71-94 (1974). https://doi.org/10.1093/genetics/77.1.71

6 Jin, E. J. & Jin, Y. A mutation linked to degt-1(ok3307) in C. elegans strain VC2633 affects rpm-1. *MicroPubl Biol* **2022** (2022). https://doi.org/10.17912/micropub.biology.000565

7 Ihara, S. *et al.* Basement membrane sliding and targeted adhesion remodels tissue boundaries during uterine-vulval attachment in Caenorhabditis elegans. *Nat Cell Biol* **13**, 641-651 (2011). https://doi.org/10.1038/ncb2233

8 Sibley, M. H., Graham, P. L., von Mende, N. & Kramer, J. M. Mutations in the alpha 2(IV) basement membrane collagen gene of Caenorhabditis elegans produce phenotypes of differing severities. *EMBO J* **13**, 3278-3285 (1994). https://doi.org/10.1002/j.1460-2075.1994.tb06629.x

9 Morck, C., Rauthan, M., Wagberg, F. & Pilon, M. pha-2 encodes the C. elegans ortholog of the homeodomain protein HEX and is required for the formation of the

pharyngeal isthmus. *Dev Biol* **272**, 403-418 (2004). https://doi.org/10.1016/j.ydbio.2004.05.011
10 Dokshin, G. A., Ghanta, K. S., Piscopo, K. M. & Mello, C. C. Robust Genome Editing with Short Single-Stranded and Long, Partially Single-Stranded DNA Donors in Caenorhabditis elegans. *Genetics* **210**, 781-787 (2018). https://doi.org/10.1534/genetics.118.301532
11 Escorcia, W., Ruter, D. L., Nhan, J. & Curran, S. P. Quantification of Lipid Abundance and Evaluation of Lipid Distribution in Caenorhabditis elegans by Nile Red and Oil Red O Staining. *J Vis Exp* (2018). https://doi.org/10.3791/57352
12 O'Rourke, E. J., Soukas, A. A., Carr, C. E. & Ruvkun, G. C. elegans major fats are stored in vesicles distinct from lysosome-related organelles. *Cell Metab* **10**, 430-435 (2009). https://doi.org/10.1016/j.cmet.2009.10.002
13 Labrousse, A., Chauvet, S., Couillault, C., Kurz, C. L. & Ewbank, J. J. Caenorhabditis elegans is a model host for Salmonella typhimurium. *Curr Biol* **10**, 1543-1545 (2000). https://doi.org/10.1016/s0960-9822(00)00833-2
14 Zhao, Y. *et al.* Two forms of death in ageing Caenorhabditis elegans. *Nat Commun* **8**, 15458 (2017). https://doi.org/10.1038/ncomms15458
